# MedDiC: high dimensional mediation analysis via difference in coefficients

**DOI:** 10.1101/2022.09.08.507169

**Authors:** Qi Zhang, Zhikai Yang, Jinliang Yang

## Abstract

High dimensional mediation analysis has been receiving increasing popularity, largely motivated by the scientific problems in genomics and biomedical imaging. Previous literature has primarily focused on mediator selection for high dimensional mediators. In this paper, we aim at the estimation and inference of overall indirect effect for high dimensional exposures and high dimensional mediators. We propose MedDiC, a novel debiased estimator of the high dimensional overall indirect effect based on difference-in-coefficients approach. We evaluate the proposed method using intensive simulations and find that MedDiC provides valid inference and offers higher power and shorter computing time than the competitors for both low dimensional and high dimensional exposures. We also apply MedDiC to a mouse f2 dataset for diabetes study and a dataset composed of diverse maize inbred lines for flowering time, and show that MedDiC yields more biologically meaningful gene lists, and the results are reproduciable across analyses using different measures of identical biological signal or related phenotype as the outcome.

Upon the acceptance of the paper, the code will be available on GitHub (https://github.com/QiZhangStat/MedDiC).

## 1 Introduction

Causal mediation analysis has been applied to many fields, such as genomics, medical imaging, epidemiology and psychology [3, 13, 26, 14, 46]. It partitions the total effect (TE) of the exposures (**Z**, Figure 1a) on the outcome (**Y**) into two parts, the indirect effect (IE, or mediation effect) that is through some measured mediating variables (**M**), and the direct effect (DE) through other mechanisms.

**Figure 1:**
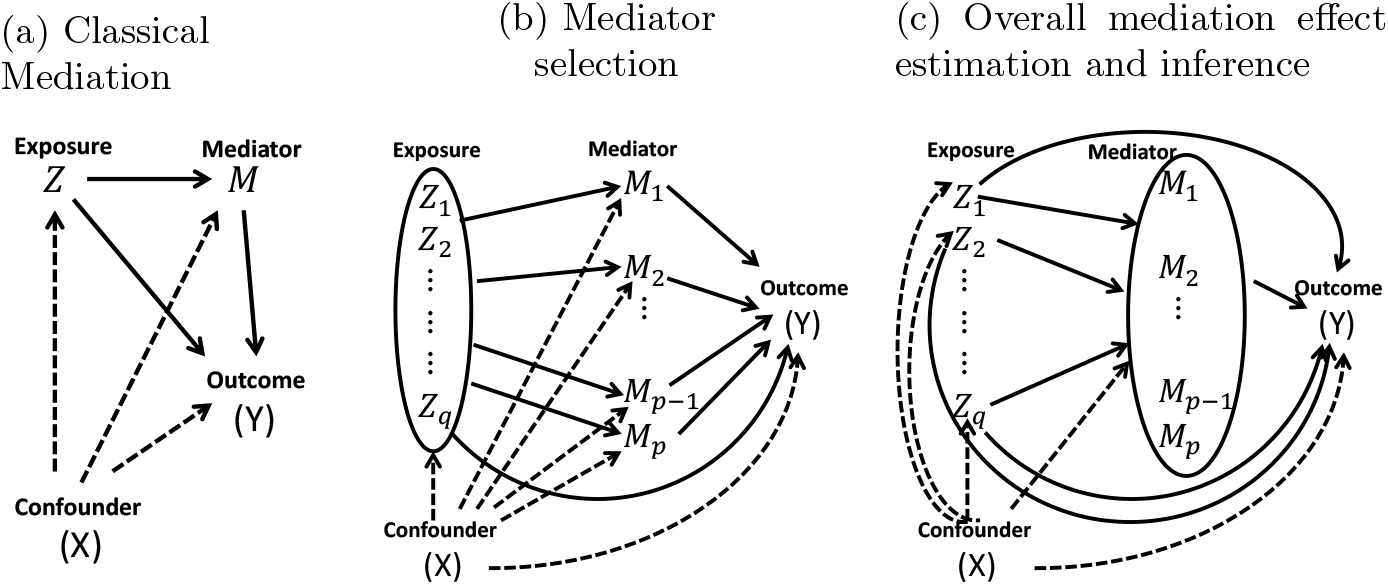
Causal diagrams for different mediation problems: classical mediation analysis (Left), mediator selection (Middle) and estimating the overall mediation effect (Right).

The classical mediation analysis studies the mediation effect of one exposure through one mediator (Figure 1a; See [1] for a recent evaluation in the context of GWAS). During the last decade, mediation analysis for multivariate low dimensional mediator have also been well studied in the literature [37, 38, 4, 28]. Largely motivated by the scientific problems in genomics and biomedical imaging, mediation models for high dimensional mediators have began drawing more and more attentions in statistics and bioinformatics. The majority of the literature in high dimensional mediation have studied mediator selection, i.e., identifying the mediators with significant mediation effect. These works usually assume that the mediators are not causally related (Figure 1b). [44] proposed HIMA, a high dimensional mediator selection method for a single exposure. It is based on intersection-union tests after screening and penalization-based estimation. Many novel methods with similar rationale have been developed for various high dimensional mediation problems, such as [43] for survival outcome, [8] for testing the significance of a group of mediators with potentially non-linear mediation effect, and [49, 48, 45] for the aggregated indirect effects of the individual mediators when the exposures are potentially high dimensional. [31, 30, 29] proposed Bayesian solutions for high dimensional mediator selection problems. Somewhat related, [3, 12, 47] introduced PCA-based approaches for high dimensional mediation analysis that essentially assume the same diagram among the mediators in Figure 1b after a data-driven linear transformation of the mediators. When the mediation effect may differ among sub-populations, [41] developed a penalization-based heterogeneous mediation analysis framework for univariate exposure and high dimensional mediator when the direct and indirect effects may vary across subgroups. [50] studied a different high dimensional mediation problem. For low dimensional exposures, they proposed Freebird, a procedure for the estimation and inference of the overall indirect effect of each exposure on the outcome through the high dimensional mediators (Figure 1c).

In this paper, we investigate the estimation and inference of the overall indirect effect through high dimensional mediators for high dimensional exposures. Our inference target is similar to [50], but we aim at high dimensional exposures. [50] does not generalize to the high dimensional exposures because it uses ordinary least squares for estimating the association between the exposures and the mediators, and their target parameter for the indirect effect is a high dimensional linear combination of the regression coefficients. We take a completely different perspective and develop **Med**iation analysis via **D**ifference **i**n **C**oefficients (**MedDiC**), a novel estimation and inference procedure for the indirect effect of high dimensional exposure through high dimensional mediators based on the difference-in-coefficients approach [21, 22].

The remaining of the paper are organized as the following. In Section 2, we present the model setup for our high dimensional mediation problem, and derive **MedDiC** estimator and its asymptotic distribution. We then present our simulation results in Section 3, the analysis of a mouse dataset and a maize dataset in Section 4, and close the paper with discussions in Section 5.

## 2 Mediation analysis with high dimensional exposure and high dimensional mediators

For *n* subjects from the population, let *Y* be the length *n* response vector, *Z* the *n* × *q* exposure matrix, *X* the *n* × *s* confounder matrix including the intercept, *M* the *n* × *p* mediator matrix. We consider the following linear mediation model

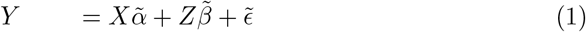

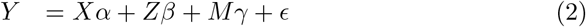

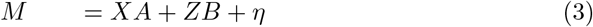

where *β* and 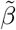 are vectors of length *q*, α and 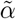 are vectors of length *s, γ* is a vector of length *p, A* and *B* are *s* × *p* and *q* × *p* matrices, respectively, 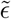, *ϵ* are length *n* noise vectors, and *η* is *n* × *p* noise matrix.

If the linear model assumption is true, there are

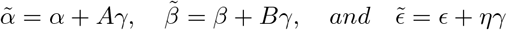

Hence only either (1)-(2) or (2)-(3) are needed for mediation analysis.

The literature of mediation analysis has been primarily interested in the estimation and inference of the indirect effect. If (2)-(3) are used, the parameter representing the indirect effect is *Bγ*. Since it is the product of two coefficients, it is called the product-of-coefficients approach. If (1)-(2) are considered, the indirect effect is 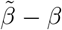, which is referred to as the difference-in-coefficients approach. The two approaches are equivalent for the above linear models, even though they target at distinct causal quantities in general [23, 28]. Which approach is preferable has been intensively discussed and evaluated in the literature without general consensus[24, 14, 21].

In the context of high dimensional mediation analysis, the product-of-coefficients approach may be necessary for mediator selection, because it enables the estimation and inference of the contributions of individual mediators. For example, HIMA [44] tests the significance of each mediator based on the joint significance test of the links *Z* → *M*_*j*_ and *M*_*j*_ → *Y*. For estimation and inference of the overall mediation effect of each elements of a low dimensional exposure, Freebird [50] also follows the product of coefficients approach to estimate *Bγ*. They plug-in *B* with its ordinary least square estimator (*Z*^*T*^ *Z*)^*−*1^(*Z*^*T*^ *M*), and then derive a debiased estimator of (*Z*^*T*^ *M*)*γ*. Since this is a high dimensional linear combination of the regression coefficients *γ*, plugging-in a debiased estimator of *γ* does not necessarily yield an unbiased asymptotically normal estimator. Thus the authors of [50] proceeded to derive their own debiased estimator. Since ordinary least square is used, it cannot be directly applied to high dimensional exposures.

For high dimensional exposures, we adopt the difference-in-coefficients approach for estimation and inference of the overall indirect effect. Our proposed procedure **MedDiC** provides a debiased estimator for the indirect effect, and the standard errors are easy to estimate from the data. The next sections detail the derivation of **MedDiC**, and then the implementation.

### 2.1 Estimating the indirect effect using the difference of debiased lasso estimators

We consider the models (1)-(2), and rewrite them as the following

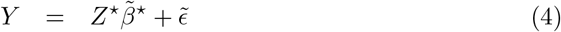

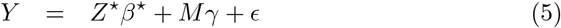

where 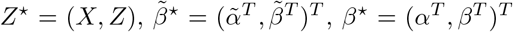. Thus the exposures are represented by the *s* + 1 to *s* + *q* columns of *Z*^⋆^. For the rest of this section, we refer to *Z*^⋆^ as the exposure matrix, because one may consider a confounder as an exposure whose effect is of no interest. We further assume that for 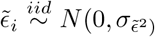 and 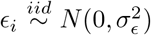. Throughout this paper, for a matrix *A*, we use *A*_ℐ, 𝒥_ to denote the submatrix of *A* such that the row and column IDs are restricted to sets ℐ and 𝒥 respectively. When ℐ = {*i*} and 𝒥 = {*j*}, we simply write it as *A*_*ij*_. We use *A*_*i*,:_ to represent the *i*th row of *A*, and *A*_:,*j*_ its *j*th column.

When the mediators and exposures are both potentially high dimensional, we advocate the use of difference-in-coefficients approach for mediation analysis. In this framework, 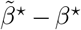 is the parameter representing the overall indirect effect for each exposure. We propose to estimate and perform statistical inference for them using 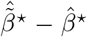 and its asymtotpic distribution, where 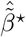 and 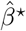 are based on the debiased lasso [42]. In the following, we derive this estimator and show that its asymptotic distribution is normal, and its variance can be easily estimated.

Based on [42], the debiased estimator of 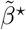 is

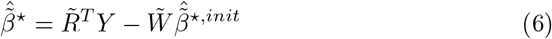

where 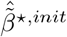 is an initial estimator of 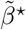, and 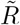 and 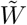 are calculated as below. Let 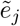 be the length *n* residual vector after regressing 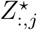, the *j*th column of *Z*^⋆^ on the other columns of *Z*^⋆^. Then 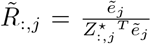 and 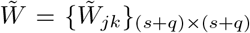 where 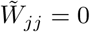 and 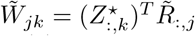 for *j* ≠ *k*.

Plugging in (4) to (6) gives

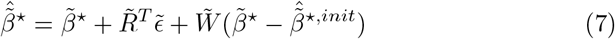

Under regularity conditions (Theorem 1 of [42]), the last term in the above is negligible and there is

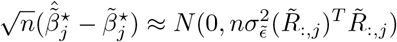

Similarly, The debiased estimator of ((*β*^⋆^)^*T*^, *γ*^*T*^)^*T*^ is

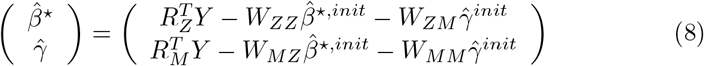

Where 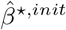 and 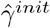 are initial estimates of *β*^⋆^ and *γ*, and *R* = (*R*_*Z*_, *R*_*M*_) and 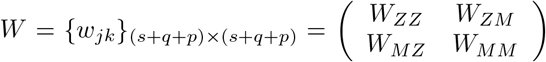 are defined in a similar fashion after substituting *Z*^⋆^ with (*Z*^⋆^, *M*). Plugging in (5) into (8), there is

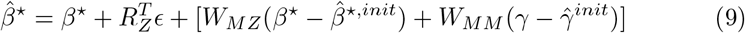

Under similar regularity conditions except the change in covariate matrix, 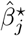 is asymptotically normal with

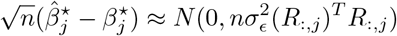

A natural estimator of 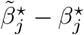 and 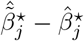. Combining (7) and (9) yields

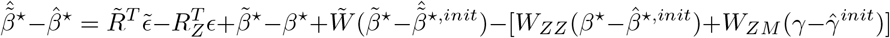

The regularity conditions of the debiased lasso [42] ensure that the two bias terms) 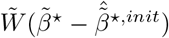 and 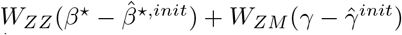 are negligible.Consequently, the bias of 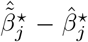 is negligible, and it is also asymptotically normal. Its variance is

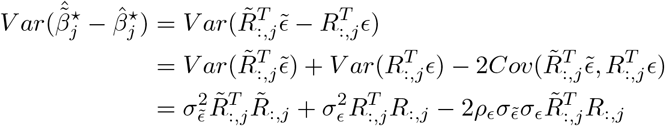

where 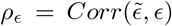. This variance formula is essentially the same as the corrected formula of McGuigan-Langholtz standard error [22] for the difference-in-coefficients estimator of the indirect effect in the case of univariate exposure and univariate mediator as presented in [21].

To summarize, if the regularity conditions of debiased lasso [42] are satisfied for both of (4) and (5), there is

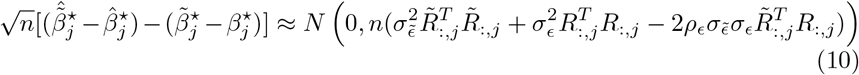

Here 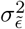 and 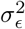 can be estimated based on the scaled lasso, and *ρ*_ϵ_ can be estimated using the empirical correlation of the residuals from the two scaled lasso regression.

The assumptions for causal interpretation of the direct and indirect effects are similar to what have been presented in the literature [50, 38]. We assume that there are no unmeasured confounding of the exposure-outcome, mediator-outcome or the exposure-mediator relationship, and the exposures do not affect any confounder between mediators and outcome.

### 2.2 Implementation

Most of the current implementations of the debiased lasso and the scaled lasso are slow or does not accommodate our mediation problem directly [5, 32, 42]. Thus we implement MedDiC from scratch using RCPP, combining the strengths of RcppArmadillo library[7] and some existing algorithms scattered in various packages such as fastGGM [40] and hdi[5].

To reduce bias in the initial estimator, we uses a scaled adaptive lasso (Algorithm 1) as the initial estimator for the debiased lasso. It uses the estimated coefficients from an initial fit to construct the weights for adaptive lasso. Similar strategies of constructing weights for adaptive lasso have been used in the literature [45, 51]. The noise level is estimated using least square after the scaled lasso, because [25] shows that it is less biased than the direct output of the original scaled lasso. We remark that [25] also advocates using cross-validation for noise level estimation, and [5] suggests to use lasso with cross-validation for the initial estimator of the debiased lasso. However, we have found no clear gain in high dimensional setting that justifies the additional computational cost of cross-validations in our exploratory simulation experiments. The score calculations are also based on the scaled lasso. The tuning parameter of the scaled lasso is set to be 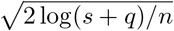 for the models without the mediators, and 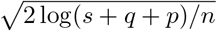 for models with mediators. For the low dimensional problems with *q* + *s < c n* where *c∈* (0, 1) is a fixed constant, the ordinary least square is used instead of the debiased lasso for estimating 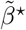. In this study, we set *c* = 0.2.

#### Algorithm 1

Scaled adaptive lasso for linear model *y* ∼ *N* (*Xβ, σ*^2^*I*_*n*_)

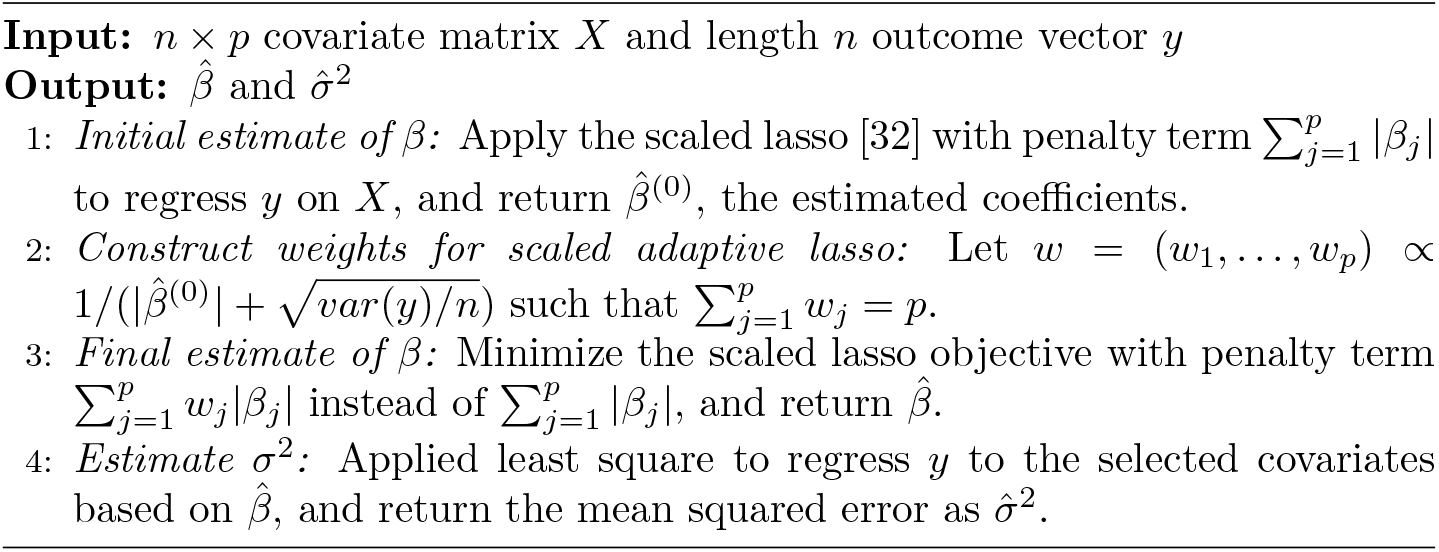

The proposed MedDiC procedure is summarized as Algrorithm 2.

#### Algorithm 2

MedDiC: Mediation analysis using Difference in Coefficients

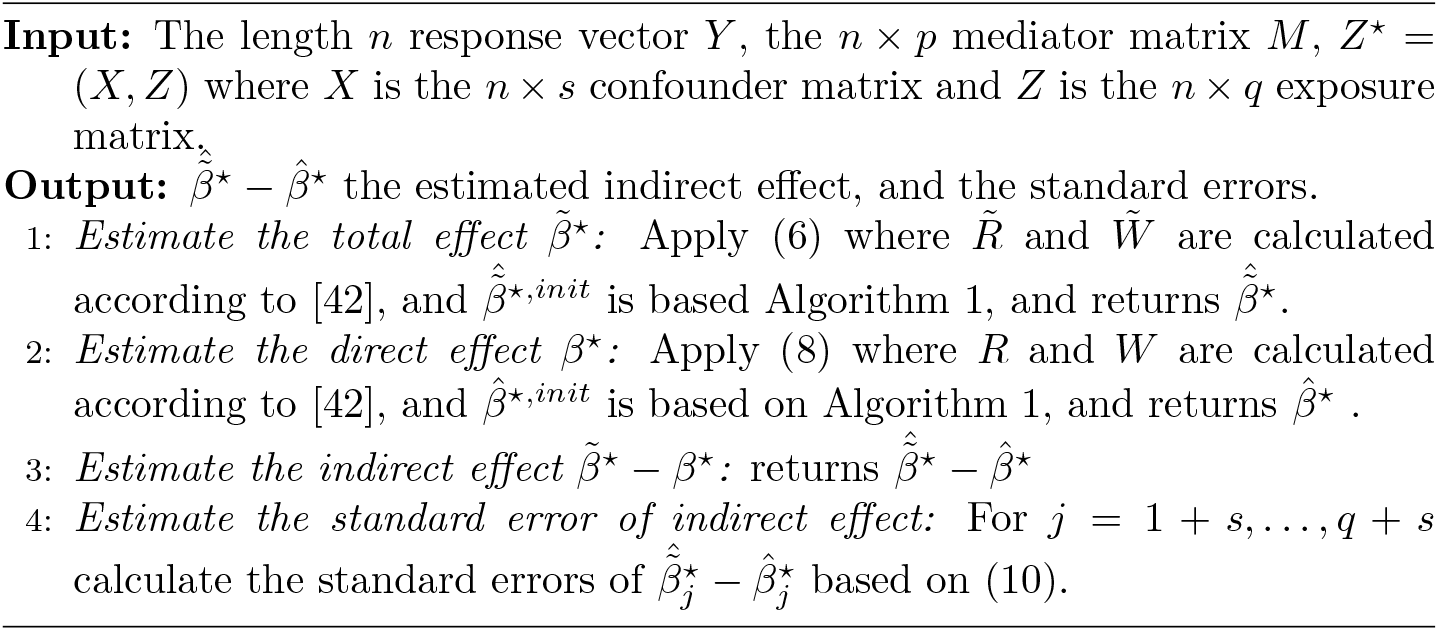

## 3 Simulation Results

### 3.1 Simulation model

We simulate the data from the conventional linear mediation model (2)-(3). For *i* = 1, …, *n*, we simulate *X*_*i*,:_ = (*X*_*i*1_, …, *X*_*is*_) ∼ *N*(0, *I*_*s*_), *Z*_*i*,:_ = (*Z*_*i*1_, …, *Z*_*iq*_) ∼*N* (0, Σ) where 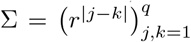 is a toeplitz matrix, *η*_i_ ∼ *N* (0, *I*_*p*_) and ϵ_*i*_ ∼*N* (0, 1). We fix *s* = 2, *α* = (−0.2, 0.2)^*T*^, and *A* is a matrix whose rows are random permutations of (−0.1, 0.1).

Recall that the indirect and the direct effect of exposure *j* are (*B*_*j*,:_*γ, β*_*j*_). We simulate *β, B* and *γ* such that randomly selected four out of *q* exposures have non-zero indirect and/or direct effects. The effects (*B*_*j*,:_*γ, β*_*j*_) of these four exposures are (−*τ*, 0), (0, − *τ*), (*τ, τ*), and (1.2*τ*, − 0.8*τ*), respectively, where *τ* is the simulation parameter for the signal strength. We remark that these values of (*B*_*j*,:_*γ, β*_*j*_) are chosen to represent the exposures with only indirect effect (complete mediation), only direct effect, both effects with the same sign, and both effects with different signs.

In the following, we describe how *γ* and *B* are simulated to satisfy the above requirement. Let *p*_*m*_ be the number of true mediators. We first simulate *γ* as a sparse vector with 2*p*_*m*_ non-zero elements with value 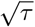. This choice is made so that the non-zero elements in both *B* and *γ* grow as the signal strength *τ* increases. Half of these 2*p*_*m*_ candidate mediators are true mediators, and we refer the other *p*_*m*_ as “fake mediators”. They have non-zero coefficients in the outcome model, but zero link to the exposures. Let *C*_*true*_, *C*_*fake*_ ⊂ {1, …, *p*} be their coordinate sets. We initially simulate *B* as a *q* × *p* sparse matrix with 10 nonzero elements in each row. These non-zero elements are random samples from a uniform distribution between -1 and 1, and their locations are also random. Then we modify *B* by setting the elements in sub-matrices 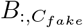 and 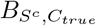 to 0, and fill 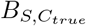 row-wise with repeating sequence of 1,0,2. The sequence of 1, 0, 2 is selected to allow each exposure with indirect effect to influence a different set of true mediators. When *p*_*m*_ = 1, however, all exposures with non-zero mediation effect must influence this only true mediator, and 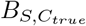 filled with the sequence of 1,1,2 instead. We further normalize *B* row-wise so that the nonzero elements of *Bγ* are − *τ, τ*, and 1.2*τ*. We then simulate *β* as a sparse vector with three nonzero elements −*τ, τ*, − 0.8*τ*. Finally, *Bγ* and *β* vectors are jointly shuffled such that there are only four exposures that have nonzero direct and/or indirect effect, and their values are as specified in the last paragraph.

In this simulation study, we fix *n* = 300,*p* = 500, *s* = 2, and consider various signal strength *τ* ∈ {0.25, 0.5, 0.75, 1, 1.5, 2}. We evaluate the performance of the proposed method for both low dimensional exposures *q* = 5, and high dimensional exposures *q* = 400. We also consider both uncorrelated exposures *r* = 0, and correlated exposures *r* = 0.4. For low dimensional exposures, we consider *p*_*m*_ = 1, 5. For high dimensional exposures, only *p*_*m*_ = 5 is considered. Each simulation setting is repeated for 25 replicates. A larger number of replicates is not used due to the high computational cost of the other methods to be compared (See Section 3.6 for a comparison of the computational times).

### 3.2 Methods to be compared

MedDiC is designed for high dimensional exposures. But it can also be applied to low dimensional exposures. Freebird [50] is the state-of-arts method for the inference of indirect effect with low dimensional exposures and high dimensional mediators. We compare MedDiC and Freebird in terms of empirical FDR, power, and the marginal coverage probability of 95% confidence intervals for low dimensional exposures.

When the exposures are high dimensional, Freebird cannot be directly applied, and there is no method for this setting except the proposed MedDiC. For benchmarking purpose, we adapt Freebird in the following two different ways for high dimensional exposures, and compare both with MedDiC.

1. Freebird After Marginal Screening (**Freebird AMS**): We first apply screening by marginal correlation between the exposures and the outcome, keep the top 20, and then apply Freebird to the survived exposures. The exposures that are not used in the final Freebird fit returns p-value= 1, and a confidence interval [0, 0]. If the true indirect effect is 0, we consider this interval cover the true value in our evaluations.
2. Freebird Univariate Scanning (**Freebird US**): This is essentially what [50] has used in their real data analysis. Let *X*_0_ be the low dimensional confounder matrix, and *U* be the top left singular vectors of the high dimensional exposure matrix *Z* (top five in simulations, and top 20 in real data analysis). For *j* = 1, …, *q*, Freebird is applied to exposure matrix (*X*_0_, *U, Z*_:,*j*_). It requires to run Freebird *q* times for a dataset. We apply this version of Freebird in our real data analysis. For simulation studies, we further modify it to reduce the computational cost. In our simulations, we only run Freebird US to a subset of exposures *S*_*o*_ ⊂{1, …, *q*}, which include all exposures with non-zero effects. This subset is chosen based on the simulation model as the following. If *j* ∈ {1, …, *q*} is an exposure with non-zero indirect or direct effect, then {*j* − 1, *j, j* + 1} ∩ { 1, …, *q*} ⊂ *S*_0_. Since there are four exposures with non-zero effects in the simulation model, Freebird US only need to run at most 12 times instead of *q* = 400 times for each simulation replicate. For exposures *j* ∉ *S*_*o*_, Freebird US returns p-value= 1, and [0, 0] as confidence interval. We remark that all true signals are used in fitting the Freebird model of Freebird US, which is different from Freebird AMS. Since the computational cost of Freebird increases roughly linearly with *q*, this modifiction allows us to include Freebird US in simulation-based comparisons at 3% of the computational cost without losing any true signals.

HIMA[44] is one of the most widely recognized mediator selection algorithm. Its statistical inference objective is different from ours, and cannot be compared with MedDiC or Freebird in general settings. As discussed in [50], HIMA’s objective is similar to that of MedDIC and Freebird only when there is only one true mediator. For the low dimensional exposure (*q* = 5) and when *p*_*m*_ = 1, we compare HIMA with MedDIC in terms of FDR and power. HIMA is designed for univariate exposure. Thus we apply HIMA to each exposure separately. We remark that *q* HIMA runs are needed for each simulation replicate. For each exposure, we use the p-value of its most significant mediator as the p-value of the overall mediation effect of this exposure.

For all methods, we adjust for multiple testing using Benjamini-Hochberg procedure[2] at level *α* = 0.05. At this level, we compare the empirical FDR and empirical power as defined in [45, 5], and also the empirical marginal coverage probability for 95% confidence intervals without adjusting for multiplicity. We plot the marginal coverage probabilities of the confidence intervals for exposures with zero and non-zero effects in separate figures, because their behaviors are different in some settings.

### 3.3 Inference of indirect effects for low dimensional exposures and high dimensional mediators

Figure 2 reports the empirical FDR and power of detecting significant indirect effects for low dimensional exposure, and when there is only one true mediator. We find that HIMA always have the highest power. However, it does not control FDR close to the nominal level when the exposures are correlated. In contrast, MedDiC and Freebird control FDR well regardless whether the exposures are independent or correlated. In the same figure, we also find that MedDiC has higher power than Freebird, especially when the signals are weak.

**Figure 2:**
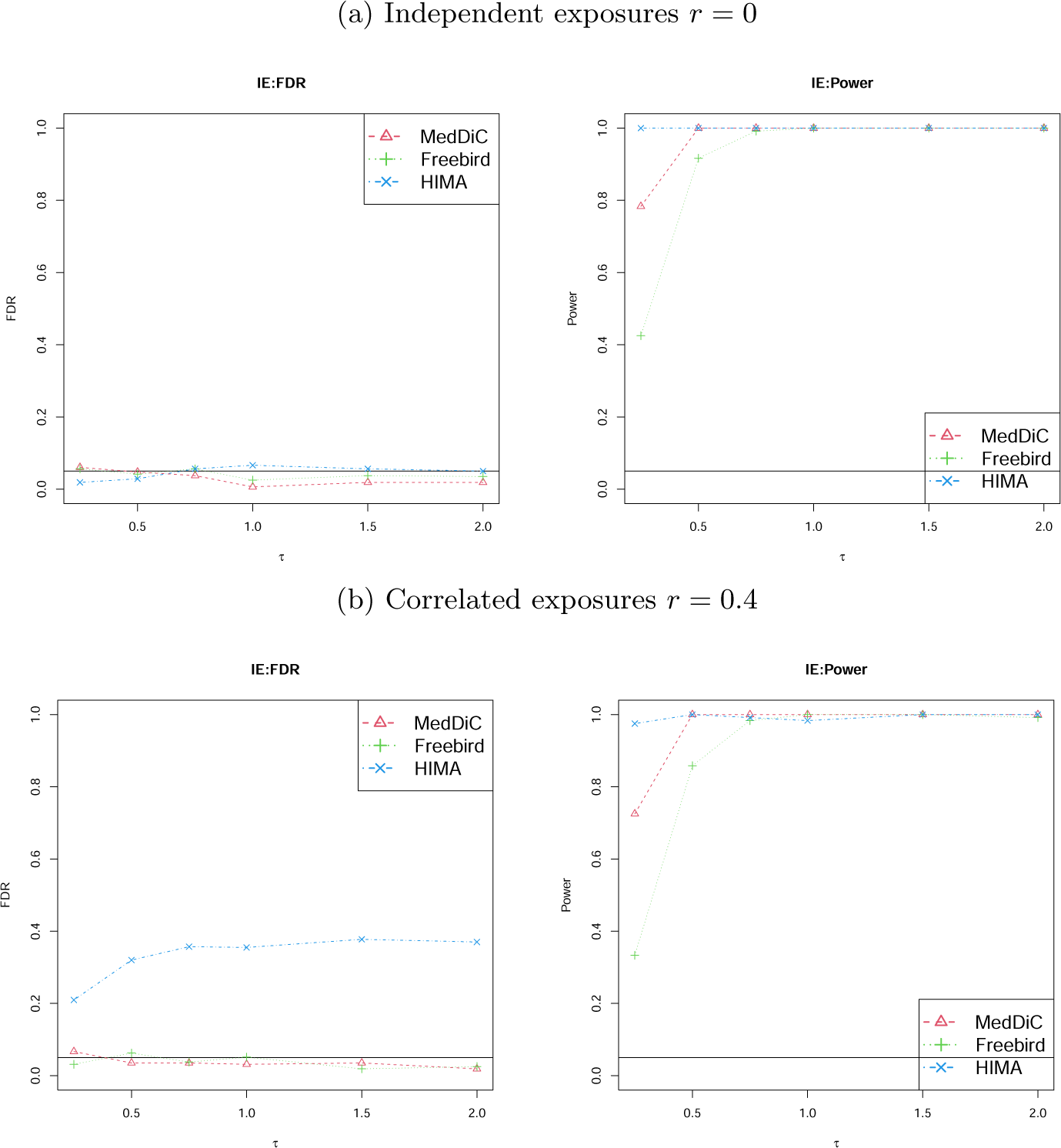
Empirical FDR and Power of detecting exposures with IE (indirect effect) when the exposure is low dimensional *q* = 5, and there is only one true mediators among the *p* = 500 candidate mediators that are included in the model. The exposures are independent (*r* = 0) in the upper panels, and correlated (*r* = 0.4) in the lower panels.

Since HIMA does not provide a confidence interval for the overall mediation effect of each exposure, we compare only MedDiC with Freebird in terms of marginal coverage probabilities in Figure 3. When the exposure is low dimensional and there is only one true mediator, we find that the coverage probabilities of MedDiC and Freebird are both close to the nominal level.

**Figure 3:**
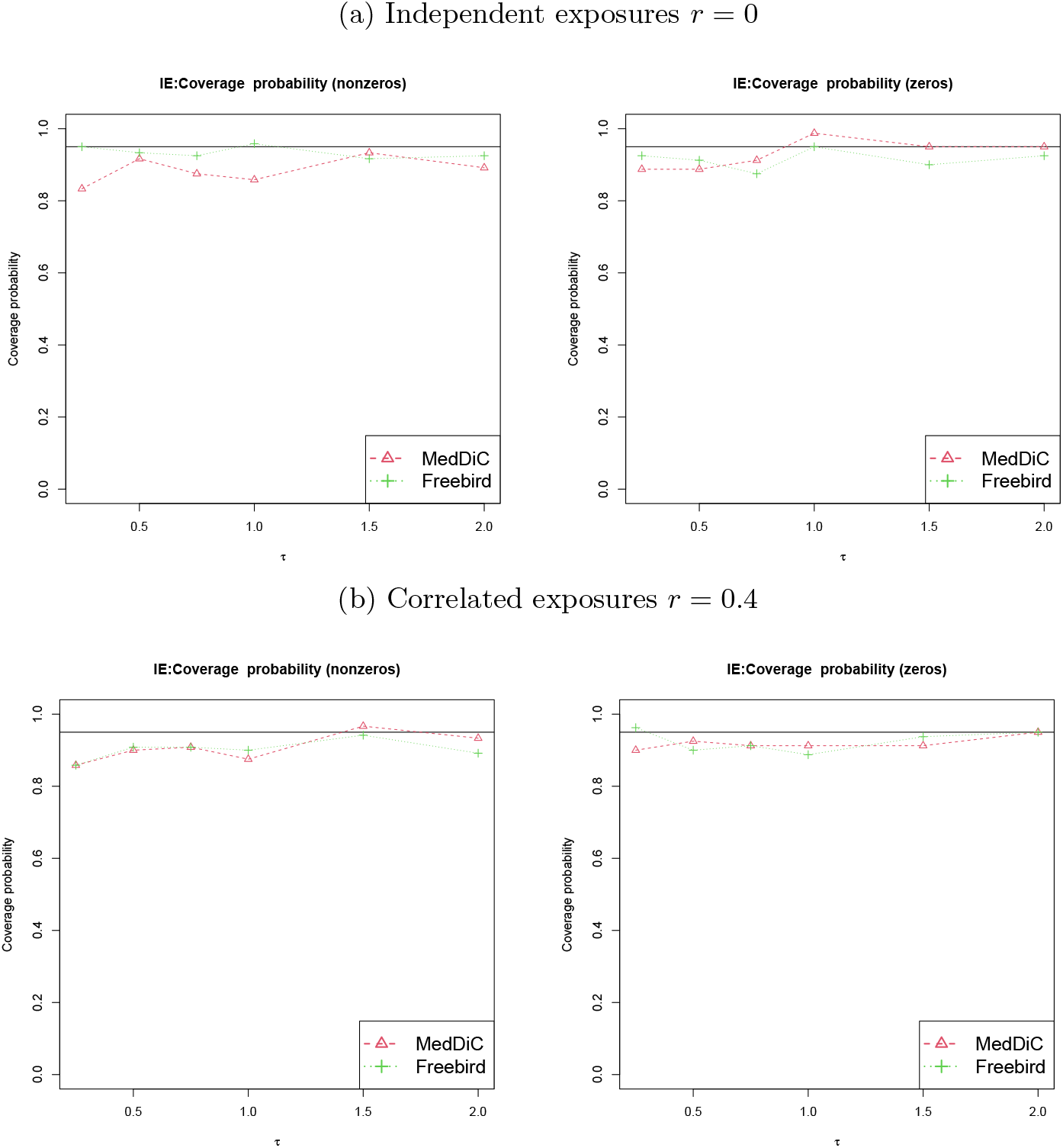
Empirical marginal coverage probabilities for the 95% confidence intervals of the IE (independent effect) when the exposure is low dimensional *q* = 5, and there is only one true mediators among the *p* = 500 candidate mediators that are included in the model. We plot the coverage probabilities for the exposures with non-zero effect and for the exposures with zero effect separately, because their behaviors may be different in some settings. The exposures are independent (*r* = 0) in the upper panels, and correlated (*r* = 0.4) in the lower panels.

In real data applications, there are usually more than one true mediator. We compare MedDiC with Freebird in the setting of low dimensional exposures (*q* = 5), and there are *p*_*m*_ = 5 true mediators among *p* = 500 potential mediators. What we observe (Figure S1-S2) confirms that MedDiC and Freebird are generally comparable for low dimensional exposures, and MedDiC has slightly higher power than Freebird for weak signals.

### 3.4 Inference of indirect effects for high dimensional exposures

When the exposures are high dimensional, MedDiC is the only method that can be directly applied. In Figure 4 and Figure S3, We compare MedDiC with Freebird AMS and Freebird US, the two adaptations of Freebird as described in Section 3.2. We find that all three methods control FDR, and the coverage probabilities of the confidence intervals for true zeros close to the nominal level. But the two adaptations of Freebird have much lower power than MedDiC. Additionally, The coverage probabilities for non-zeros from Freebird AMS are much lower than the nominal level.

**Figure 4:**
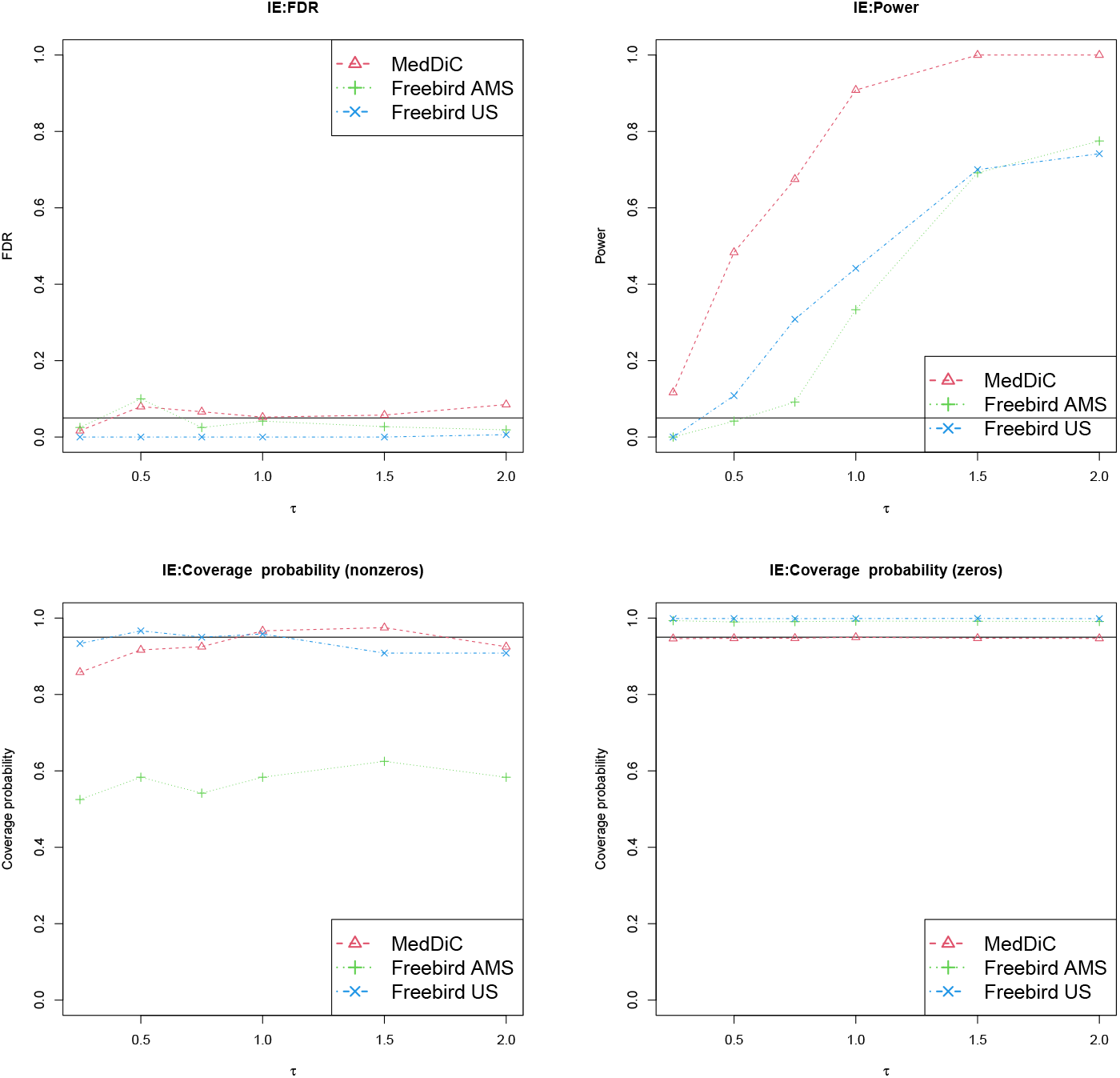
Empirical FDR (upper left) and Power (upper right) of detecting exposures with IE (indirect effect), and the marginal coverage probabilities for the true nonzero IE (lower left) and the exposures with zero IE (lower right). The exposures are high dimensional(*q* = 400), and there are five true mediators among the *p* = 500 candidate mediators that are included in the model. The exposures are independent (*r* = 0).

### 3.5 Complimentary Inference of direct effects and total effects

The major focus of this paper is the inference of indirect effect. We do not claim the inference of the direct and total effects based on MedDiC as a novel contribution, because they are the direct outputs of the debiased lasso applied to (1) and (2). Nevertheless, we report these complimentary inference results of direct effects and total effects from MedDiC in Figures S4-S9. We find that the inference of the total effect is always valid in terms of FDR control and coverage probabilities of the confidence intervals, and the power is reasonable. The statistical tests of direct effects return reasonable control of Type I error, and high power. The coverage probabilities of the non-zero direct effects are generally lower than the nominal level.

### 3.6 Computational time comparison

We also evaluate the computational cost of MedDiC and compare with Freebird and HIMA on the same computer. This is a laptop with Intel(R) i7-1065G7 CPU @1.30*GHz* 1.50*GHz*, 32 GB memory, Win 10 operation system and Microsoft R Open 4.02 for computing. In Table 1, we present the average computational time (in seconds) for the simulation setting with (*q, p*_*m*_, *r, τ*) = (5, 1, 0, 0.5). We find that the computational cost of MedDiC is less than 1% of Freebird, and less than 5% of HIMA. It is potentially partially due to its efficient implementation using RCPP. We do not document the computational time of the two high dimensional adaptations of Freebird. This is because they are expected to be much slower than Freebird with *q* = 5 as more exposures are included in the model fitting. We only present the computational time of MedDiC for (*q, p*_*m*_, *r, τ*) = (400, 5, 0, 0.5), and find that it is already smaller than the computational time of Freebird and HIMA in the low dimensional setting. We remark that we do not use a larger number of replicates in our simulation studies largely due to the high computational cost of Freebird and its adaptations.

**Table 1:** Average computational time (in seconds) of MedDiC, Freebird, and HIMA for the simulation setting (*q, p*_*m*_, *r, τ*) = (5, 1, 0, 0.5) and the computational time of MedDiC for the simulation setting (*q, p*_*m*_, *r, τ*) = (400, 5, 0, 0.5)

| $(q, p_m, r, \tau)$ | $(5, 1, 0, 0.5)$ | | | $(400, 5, 0, 0.5)$ |
| --- | --- | --- | --- | --- |
| Method | MedDiC | Freebird | HIMA | MedDiC |
| Average time | 2.83 | 365.39 | 68.115 | 29.57 |

## 4 Real data analysis

### 4.1 Dissection of a colocalized QTL peak for insulin and islet-specific eQTL hotspot on chr2 of *Mus musculus*

We analyze a mouse f2 cross data for diabetes study [39, 35, 34]. [33] identified an islet specific eQTL hotspot at around 75cM on chr2 that regulated a wide range of genes, including both cis- and trans-regulation. It is widely believed that trans-eQTLs co-mapped at a trans-eQTL hotspot are likely to be co-regulated by common regulator genes, such as transcription factors, that have cis-eQTLs at this locus. [16] applied this approach to dissect this hotspot using this dataset. Instead of all eQTLs mapped to this locus, they focused on the 129 genes with human homologues associated to T2D based on the previous literature, and attempted to explain the trans-eQTLs of these genes using their cis-eQTLs. They identified Nfatc2, a transcription factor as a candidate regulator of the majority of the 129 genes. However, [16] does not establish any link between Nfatc2 and any diabetes related phenotypes measured in this dataset.

There is also a QTL peak for insulin close to this eQTL hotspot. When eQTLs and QTL colocalize, the genes are likely to be regulating the phenotype. Combining this with the approach in [16, 34], we study the causal mediation relationship among the cis-regulated genes, trans-regulated genes, and the phenotype using high dimensional mediation framework. In this framework, the exposures are the expression of the genes with cis-eQTL in this locus, the mediators are the expression of the genes with trans-eQTL mapped to this region, the response is the phenotype, and the confounders are genetic markers. This is different from many genetical applications of high dimensional mediation in which the genetic markers are the exposures. This dataset has only 2057 SNPs across the whole mouse genome. Thus they are more useful as markers of the genetic location, and the causal effect of these individual SNPs are not biologically informative. In contrast, there are more than 40k transcripts in the gene expression dataset, and identifying the genes that initiates the regulation of insulin through the colocalization of QTL and eQTL is of the interests of the biologists as suggested in [33]. The gene expressions are measured in islet, the tissue that makes insulin. In our analysis, the exposures identified with total effect are those cis-regulated genes that are associated with insulin. The exposures identified with indirect effect are the cis-regulated genes that regulate insulin through regulating the expression of the trans-regulated genes. Consequently, they are more likely to be regulator genes such as transcription factors in a similar spirit of [16]. The exposures identified with direct effect are the cis-regulated genes that are associated with the phenotype through other mechanisms that do not involve the trans-regulated genes under consideration.

We decide to focus on a region from 120Mbps to 170Mbps on chr2, which include the center of the eQTL hotspot and the QTL peak. We define an eQTL as cis-if the distance between the transcription starting site of the gene being regulated and the genetic marker is less than 10Mbps. This leads to 193 cis-regulated genes as exposures, and 4561 trans-regulated genes as the mediators. This dataset has three independent measures of the blood insulin level at sacrifice (named “10wk”,“10wkRep”, and “rbm”), which provides us a unique opportunity to evaluate statistical methods based the reproducibility of the analysis results across independent measurements of identical biological signal. We perform three high dimensional mediation analysis using the same confounders, exposures and mediators, and each of the three measurements of insulin as the outcome. For each analysis, we remove the mouse with missing value in the particular phenotype. There were 491 mice in total in the dataset with gene expression data, the number of mice in the three analyses are 491,488 and 490, respectively. This case study is a setting in which the exposures are not high dimensional (*q < n*), but the confounders are (*s > n*). MedDiC can still be applied, but not Freebird. For Freebird, we use the top 20 singular vectors of the genotype matrix as the confounders so that 20 + *q < n*. We exclude HIMA from comparison because its objective is different as have described in the previous sections. Both methods return the results at significance level *FDR* = 0.1.

Table 2 presents the number of each type of effects in each analysis, and the overlap of the same type of effects between each pair of phenotypes. We find that MedDiC identifies more exposures with indirect effect than Freebird, while still maintaining similar level of overlaps. More than half of the indirect effects are identified in at least two analysis.

**Table 2:** Results of Mouse data: Number of cis-regulated genes selected with each type of effect from each analysis, and their overlaps across phenotypes.

|  | 10wk | 10wkRep | rbm | 10wk,<br>10wkRep | 10wk,<br>rbm | 10wkRep,<br>rbm |
| --- | --- | --- | --- | --- | --- | --- |
| Indirect<br>(MedDiC) | 40 | 46 | 44 | 30 | 21 | 24 |
| Direct<br>(MedDiC) | 2 | 3 | 5 | 2 | 1 | 1 |
| Total<br>(MedDiC) | 22 | 14 | 24 | 12 | 15 | 10 |
| Indirect<br>(Freebird) | 13 | 31 | 16 | 12 | 10 | 14 |

Next, we compare the methods based on the regulatory potential of the identified effects of each type. Recall that the exposures with the indirect effects are the cis-regulated genes that regulates the trans-eQTLs, which in turn regulate the outcome. Thus they are more likely to be regulators such as transcription factors than the exposures with direct and or total effects but no indirect effect. We utilize three lists of known transcription factors for *Mus musculus* [10, 11, 20]. Each of them have 1385[20],1374[10], and 1636[11] transcription factors listed, and they are largely consistent, with 1042 genes present in all three lists. Table 3 presents the overlap between the model outputs and each of the gene lists. The sizes of these gene lists are small relative to the total number of genes, and number of transcription factors at the locus under consideration will be even much smaller. Consequently, we do not expect large overlap sizes. Nevertheless, the contrasts in Table 3 is clear. Many indirect effects identified by MedDiC are known transcription factors, while Freebird output only identified one known transcription factor which is also identified by MedDiC. MedDiC finds no transcription factors among the exposures with no indirect effect but only direct or total effects. The transcription factors selected are fairly consistent across all three phenotypes and three databases. Especially, MGA, E2F1 and FOXa2 present in all nine combinations.

**Table 3:** Results of Mouse data: transcription factors in the selected cisregulated genes with each effect type.

| Effect type | Phenotype | Number of genes | Fantom5 [20] | Gifford lab [10] | AnimalTFDB 3.0 [11] |
| --- | --- | --- | --- | --- | --- |
| Indirect (MedDiC) | 10wk | 33 | mga,zfp106, foxa2,e2f1, dnmt3b,foxs1 | mga,foxa2, e2f1,foxs1 | mga,foxa2, e2f1,foxs1 |
|  | 10wkRep | 39 | mga,foxa2, ncoa6,e2f1, dnmt3b,foxs1 | mga,foxa2, e2f1,foxs1 | mga,foxa2, e2f1,foxs1 |
|  | rbm | 39 | mga,zfp106, ovol2,foxa2, e2f1,dnmt3b, id1,sall4 | mga,ovol2, foxa2,e2f1, sall4 | mga,myef2, ovol2,foxa2, e2f1,id1, sall4 |
| Direct/Total no Indirect (MedDiC) | 10wk | 9 | N/A | N/A | N/A |
|  | 10wkRep | 3 | N/A | N/A | N/A |
|  | rbm | 10 | N/A | N/A | N/A |
| Indirect (Freebird) | 10wk | 10 | N/A | N/A | N/A |
|  | 10wkRep | 26 | foxa2 | foxa2 | foxa2 |
|  | rbm | 11 | foxa2 | foxa2 | foxa2 |

### 4.2 Genetic mediation analysis of flowering time of *Zea Mays*

We apply genetic mediation analysis to the Maize 282 Association Panel, with the global diversity of public maize inbred lines [9]. The best linear unbiased prediction (BLUP) values of days to tassel (**DT**, days after planting until 50% of plants in the row shedding pollen), days to silk (**DS**, days after planting until 50% of plants in the row silking), growing degree days to tassel (**GDT**), and growing degree days to silk (**GDS**) are used as outcomes of this analysis. These traits are all closely related because they are different measures of the maize flowering time. The potential mediating genes used in this study are 420 top genes associated with male and female flowering and expressed in mature leaf during the day. Mature leaf is the tissue most relevant to flowering time in the dataset [17, 27]. The exposures are the 12,464 SNPs on chromosome 3 from 150 Mbps to 180 Mbps region, including upstream and downstream of a transcription factor *ZmMADS69* that is known to be associated with flowering time [19]. The confounders are the top 20 singular vectors of the whole genome.

After removing the lines with missing values in the phenotypes and/or gene expression, there are 191 maize inbred lines left. The dataset has 420 potential mediators, and 12,464 SNPs. Thus the exposure is high dimensional, and Freebird cannot be applied directly. We apply Freebird US, and add the top 20 singular vectors of the genotype matrix of the locus as additional confounders. Freebird US detects 55 significant indirect effects for DT, 7 for GDT and 0 for DS and GDS, while MedDiC discovers a few dozens of each type of effects for each trait (Table 4). This confirms that MedDiC is more powerful than Freebird US. Lacking ground truth at molecular level, we evaluate the outputs based on the overlaps of the same effect type between each pair of traits, and find that MedDiC yields results that are reproduciable across biologically similar traits.

**Table 4:** Results of Maize data: Number of SNPs selected with each type of effect and from each analysis, and their overlaps.

|  | DS | DT | GDS | GDT | DS,<br>DT | DS,<br>GDS | DS,<br>GDT | DT,<br>GDS | DT,<br>GDT | GDS,<br>GDT |
| --- | --- | --- | --- | --- | --- | --- | --- | --- | --- | --- |
| Indirect<br>(MedDiC) | 79 | 87 | 89 | 87 | 58 | 47 | 43 | 42 | 47 | 65 |
| Direct<br>(MedDiC) | 98 | 100 | 134 | 145 | 67 | 64 | 59 | 53 | 60 | 99 |
| Total<br>(MedDiC) | 72 | 91 | 54 | 104 | 58 | 43 | 53 | 43 | 67 | 48 |
| Indirect<br>(Freebird<br>US) | 0 | 55 | 0 | 7 | 0 | 0 | 0 | 0 | 0 | 7 |

## 5 Discussions

In this paper, we have proposed **Med**iation analysis via **D**ifference **i**n **C**oefficients (**MedDiC**) for estimation and inference of the overall indirect effect of the elements of high dimensional exposures. For high dimensional exposures, our simulation studies have shown that it achieves higher power than the two high dimensional adaptations of Freebird, and maintains the empirical FDR of the test and coverage probabilities of the confidence intervals close to the nominal level. For low dimensional exposures, its performance is comparable to Freebird. MedDiC also provides complimentary inference results of the direct and total effects. These inference results are valid, except for the coverage probability of the confidence intervals of the direct effects. MedDiC is computationally efficient, and at least 20 times faster than the competitors. Our analysis of a maize data and a mouse data show that MedDiC identifies more indirect effects than Freebird, and the identified effects are consistent across different measurements of the same biological signal or similar biological signals. In the mouse data analysis, we also show that the indirect effects identified by MedDiC are more likely to be transcription factors than direct/total effects from MedDiC or the indirect effects from Freebird.

We have primarily focused on the simple linear structural equations in this project. We have not considered any models with interactions between exposures and/or mediators, or models with nonlinear mediation effects. MedDiC can be directly applied to some of these settings and provide statistical inference for indirect effect based on the difference-in-coefficients approach. However, these estimands may be different from their counterparts based on the product-of-coefficients approach. Careful studies of MedDiC for any of these models and the justification of the causal interpretation of the results are beyond the scope of this study. MedDiC has utilized the debiased lasso as defined in [42] because it is convenient to implement. We are aware that there are other debiased estimators for high dimensional linear models [15, 36, 6, 18], some of which may have better theoretical properties or empirical performance in some settings. A comprehensive evaluation of these debiased estimators in the MedDiC framework is one of our future interest. In particular, we are interested in providing better inference for the direct effects. Another possible extension of the current work is to improve its resolution, and quantify the indirect effect through a small subset of the mediators instead of all of them. This may depend on the positions of the mediator subset under consideration in the causal graph among the potential mediators.

## Acknowledgements

We thank Dr. Sihai Dave Zhao and Ms Ruixuan Rachel Zhou for correspondences on their paper [50] and providing the code.

This research is supported by the startup fund from University of New Hampshire to QZ and the Agriculture and Food Research Initiative Grant number 2019-67013-29167 from the USDA National Institute of Food and Agriculture to JY.

## Supplementary section

### Supplementary Figures

**Figure S1:**
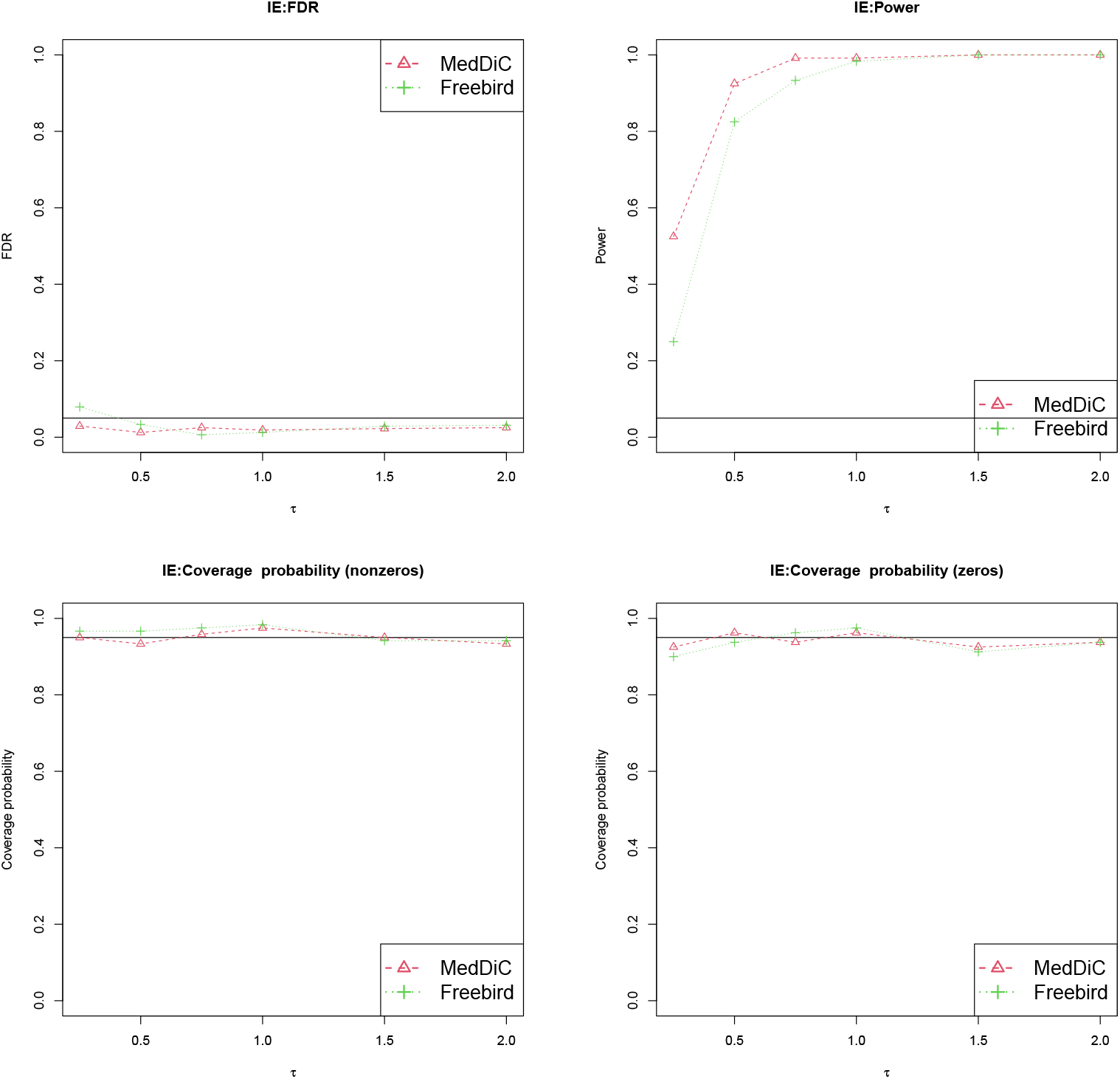
Empirical FDR (upper left) and Power (upper right) of detecting exposures with IE (indirect effect), and the marginal coverage probabilities for the true nonzero IE (lower left) and the exposures with zero IE (lower right). The exposures are low dimensional *q* = 5, and there are five true mediators among the *p* = 500 candidate mediators that are included in the model. The exposures are independent (*r* = 0).

**Figure S2:**
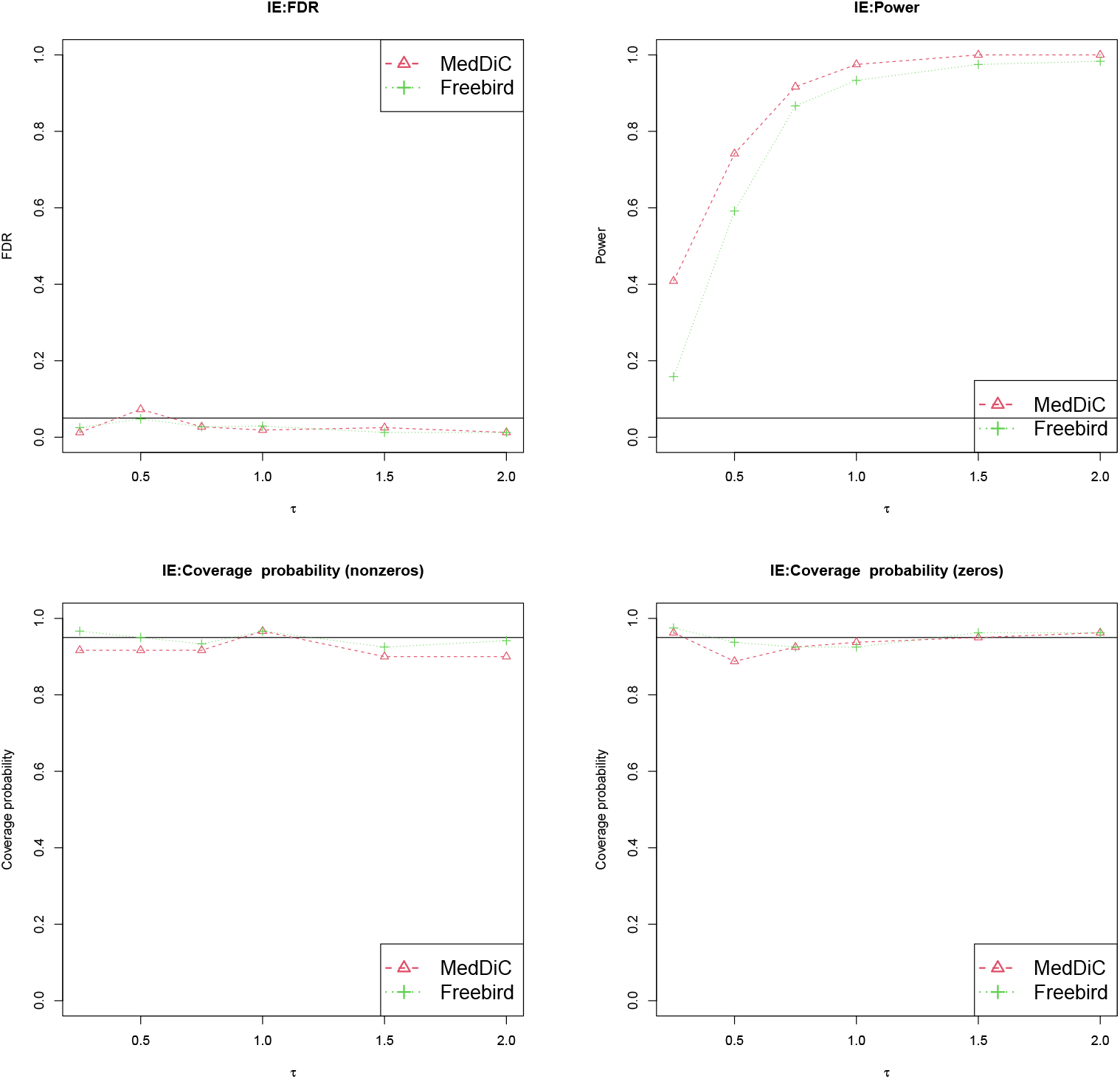
Empirical FDR (upper left) and Power (upper right) of detecting exposures with IE (indirect effect), and the marginal coverage probabilities for the true nonzero IE (lower left) and the exposures with zero IE (lower right). The exposures are low dimensional *q* = 5, and there are five true mediators among the *p* = 500 candidate mediators that are included in the model. The exposures are correlated (*r* = 0.4).

**Figure S3:**
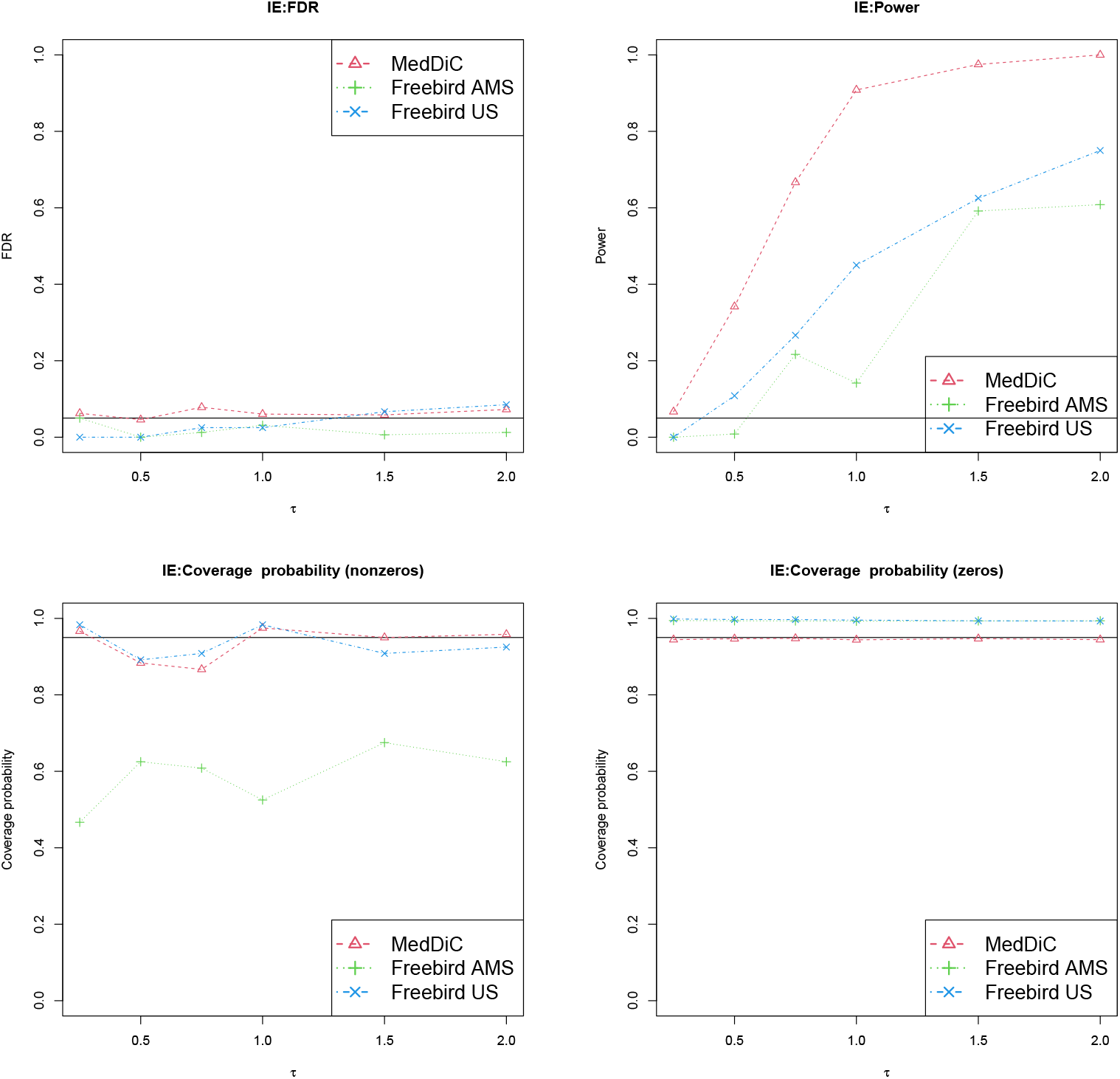
Empirical FDR (upper left) and Power (upper right) of detecting exposures with IE (indirect effect), and the marginal coverage probabilities for the true nonzero IE (lower left) and the exposures with zero IE (lower right). The exposures are high dimensional(*q* = 400), and there are five true mediators among the *p* = 500 candidate mediators that are included in the model. The exposures are correlated (*r* = 0.4).

**Figure S4:**
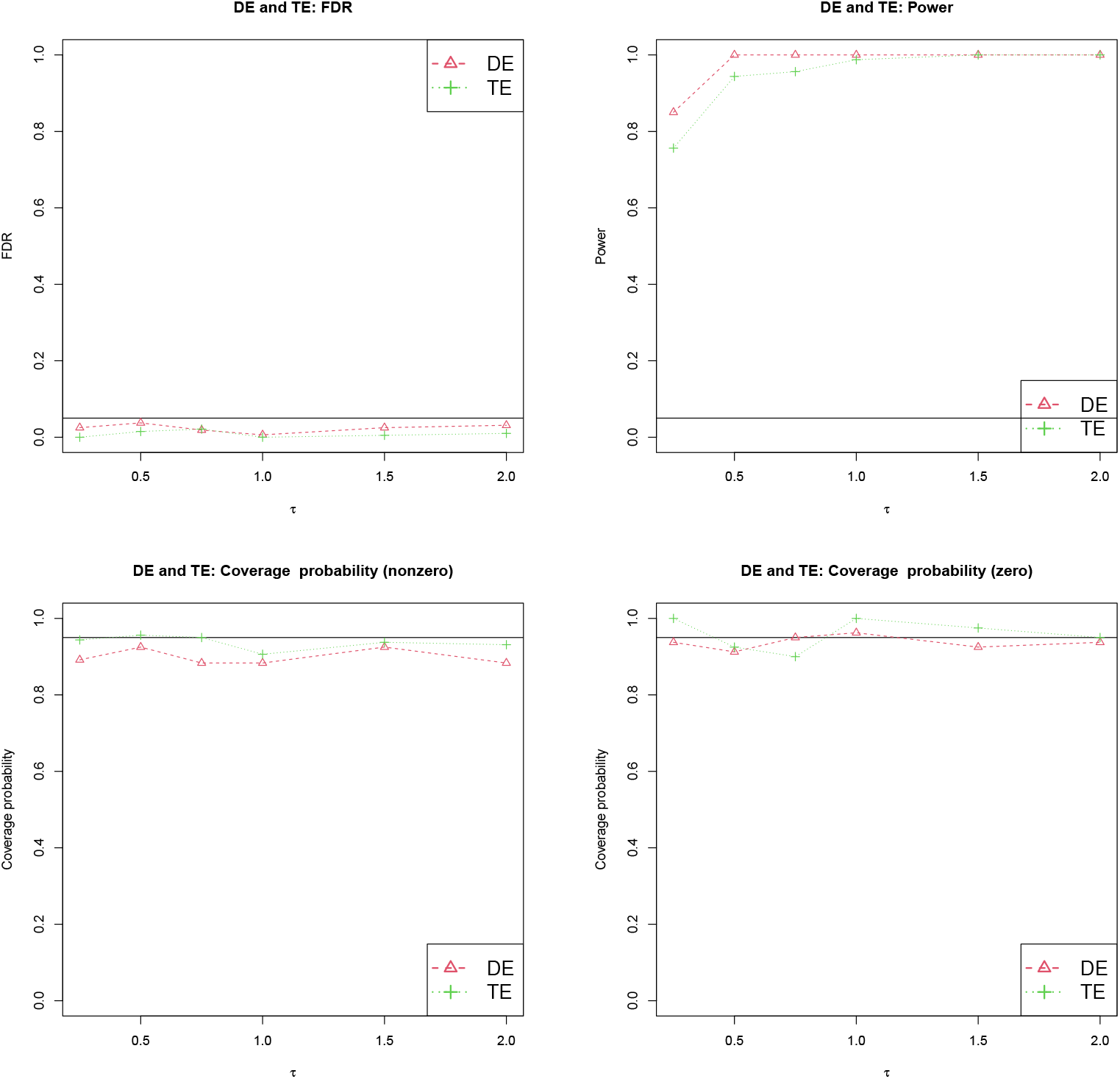
Complimentary inference of total effect (TE) and direct effect (DE) from MedDiC. Empirical FDR (upper left) and Power (upper right), and the marginal coverage probabilities for the true nonzero (lower left) and zero (lower right). The exposures are low dimensional(*q* = 5), and there is one true mediator among the *p* = 500 candidate mediators that are included in the model. The exposures are independent (*r* = 0).

**Figure S5:**
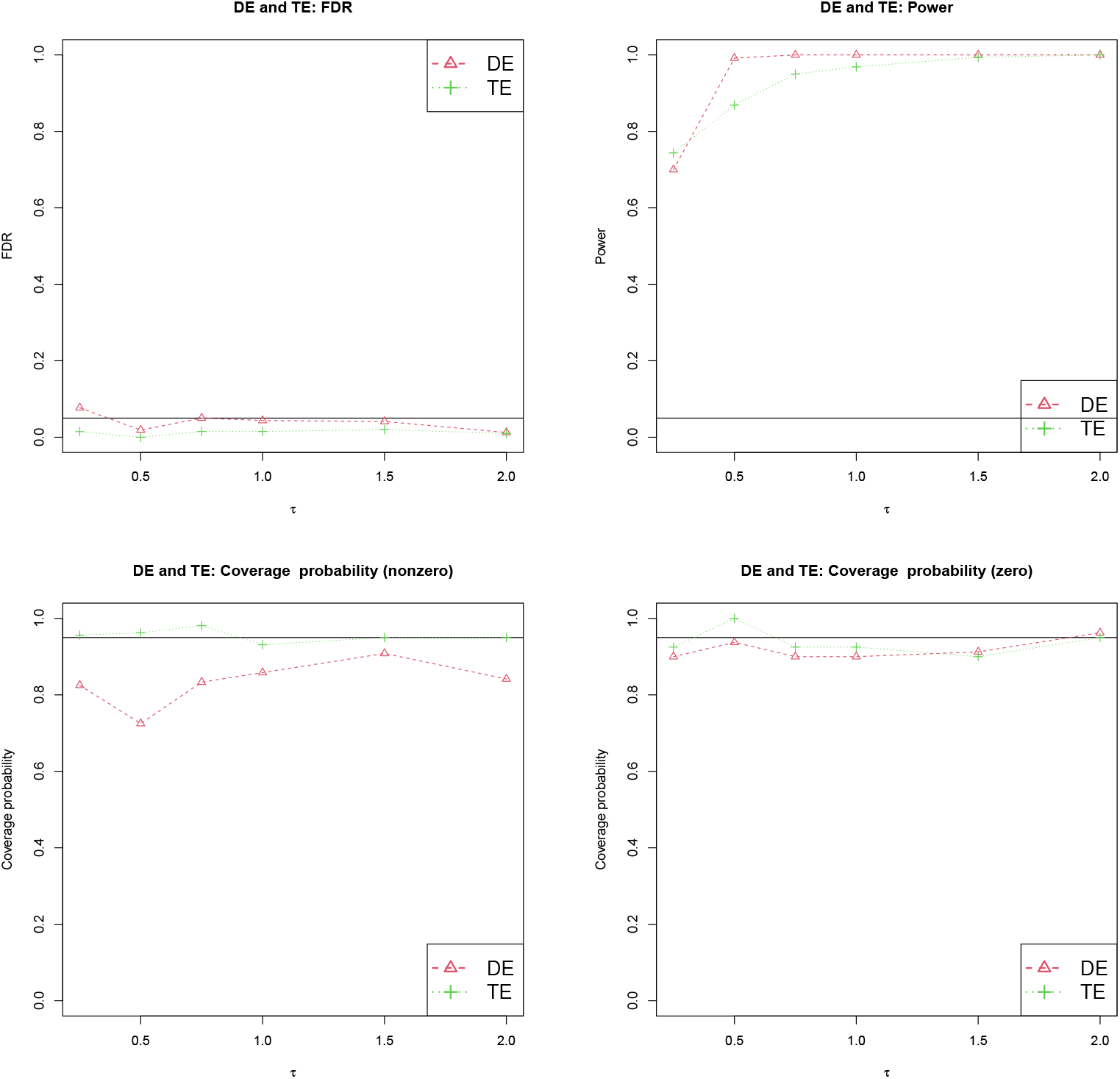
Complimentary inference of total effect (TE) and direct effect (DE) from MedDiC. Empirical FDR (upper left) and Power (upper right), and the marginal coverage probabilities for the true nonzero (lower left) and zero (lower right). The exposures are low dimensional(*q* = 5), and there is one true mediator among the *p* = 500 candidate mediators that are included in the model. The exposures are correlated (*r* = 0.4).

**Figure S6:**
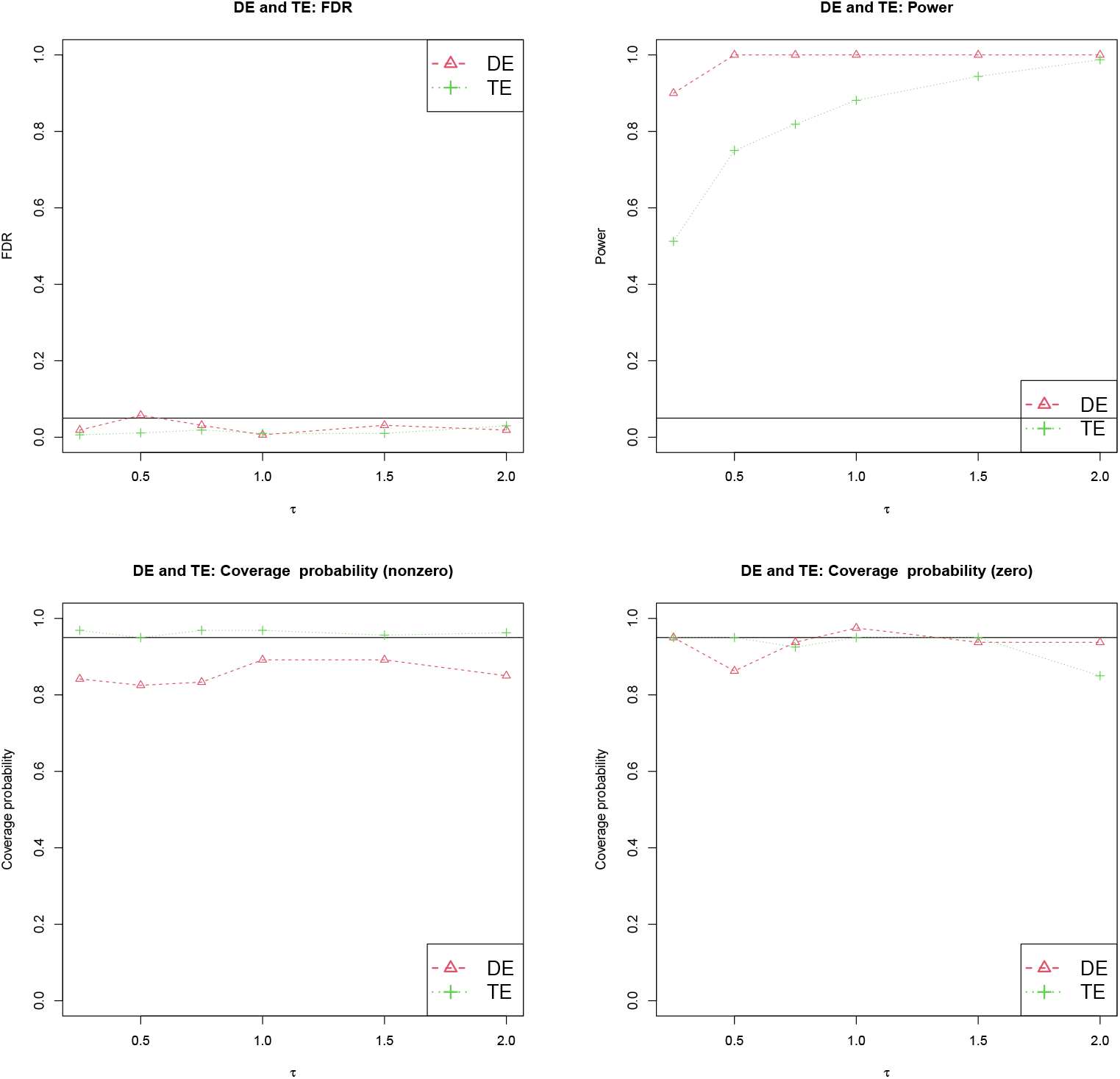
Complimentary inference of total effect (TE) and direct effect (DE) from MedDiC. Empirical FDR (upper left) and Power (upper right), and the marginal coverage probabilities for the true nonzero (lower left) and zero (lower right). The exposures are low dimensional(*q* = 5), and there are five true mediators among the *p* = 500 candidate mediators that are included in the model. The exposures are independent (*r* = 0).

**Figure S7:**
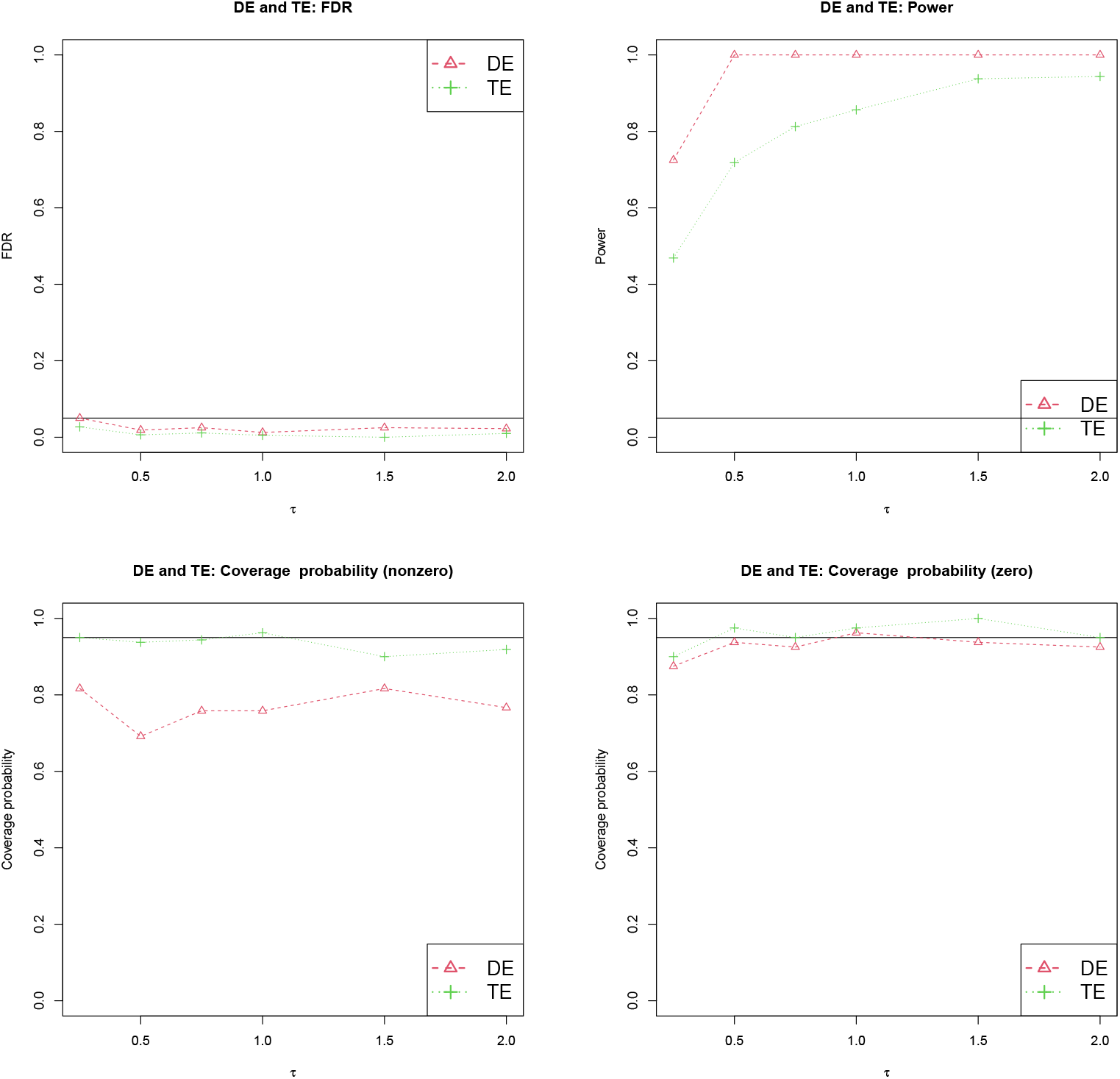
Complimentary inference of total effect (TE) and direct effect (DE) from MedDiC. Empirical FDR (upper left) and Power (upper right), and the marginal coverage probabilities for the true nonzero (lower left) and zero (lower right). The exposures are low dimensional(*q* = 5), and there are five true mediators among the *p* = 500 candidate mediators that are included in the model. The exposures are correlated (*r* = 0.4).

**Figure S8:**
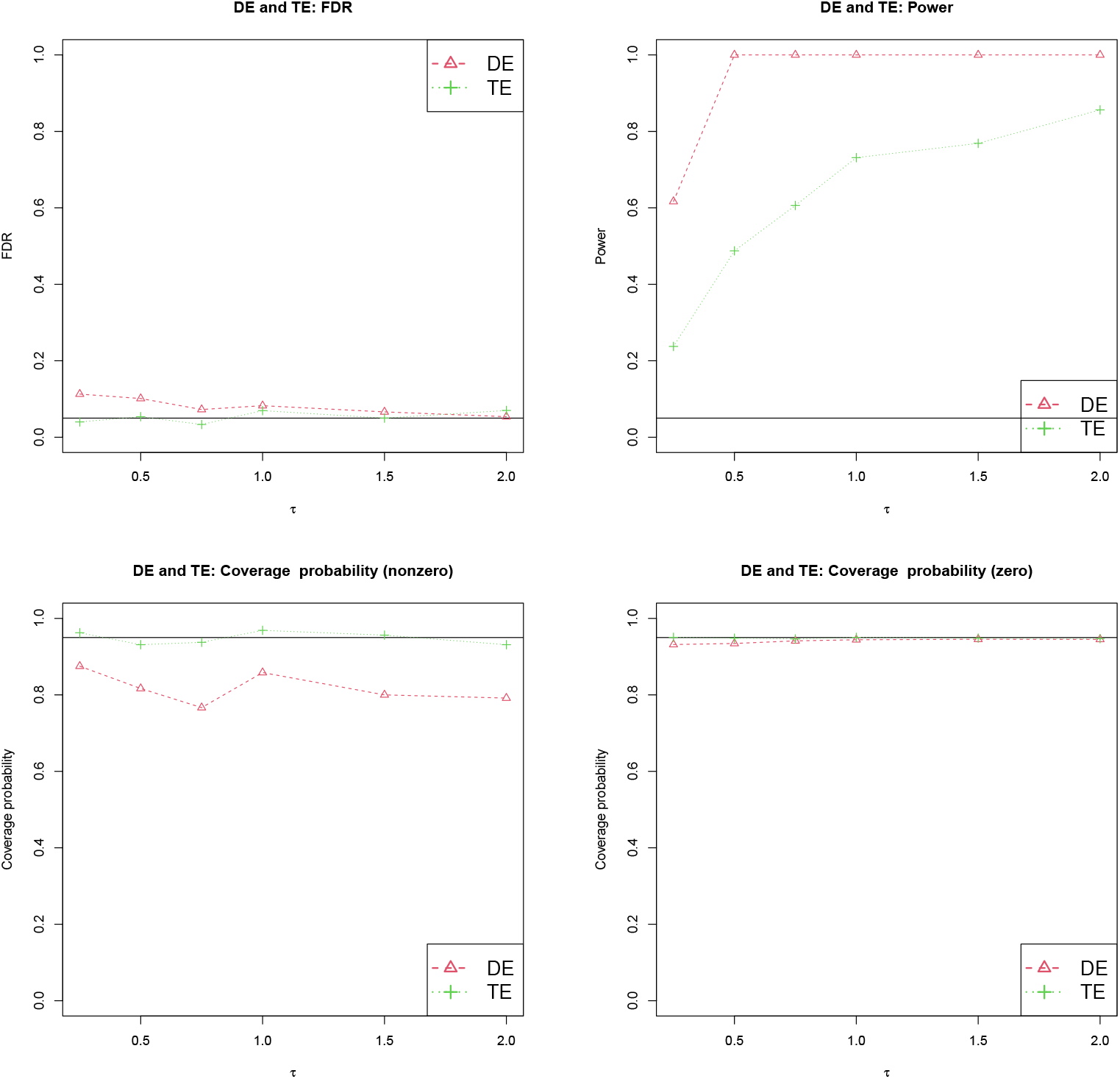
Complimentary inference of total effect (TE) and direct effect (DE) from MedDiC. Empirical FDR (upper left) and Power (upper right), and the marginal coverage probabilities for the true nonzero (lower left) and zero (lower right). The exposures are high dimensional(*q* = 400), and there are five true mediators among the *p* = 500 candidate mediators that are included in the model. The exposures are independent (*r* = 0).

**Figure S9:**
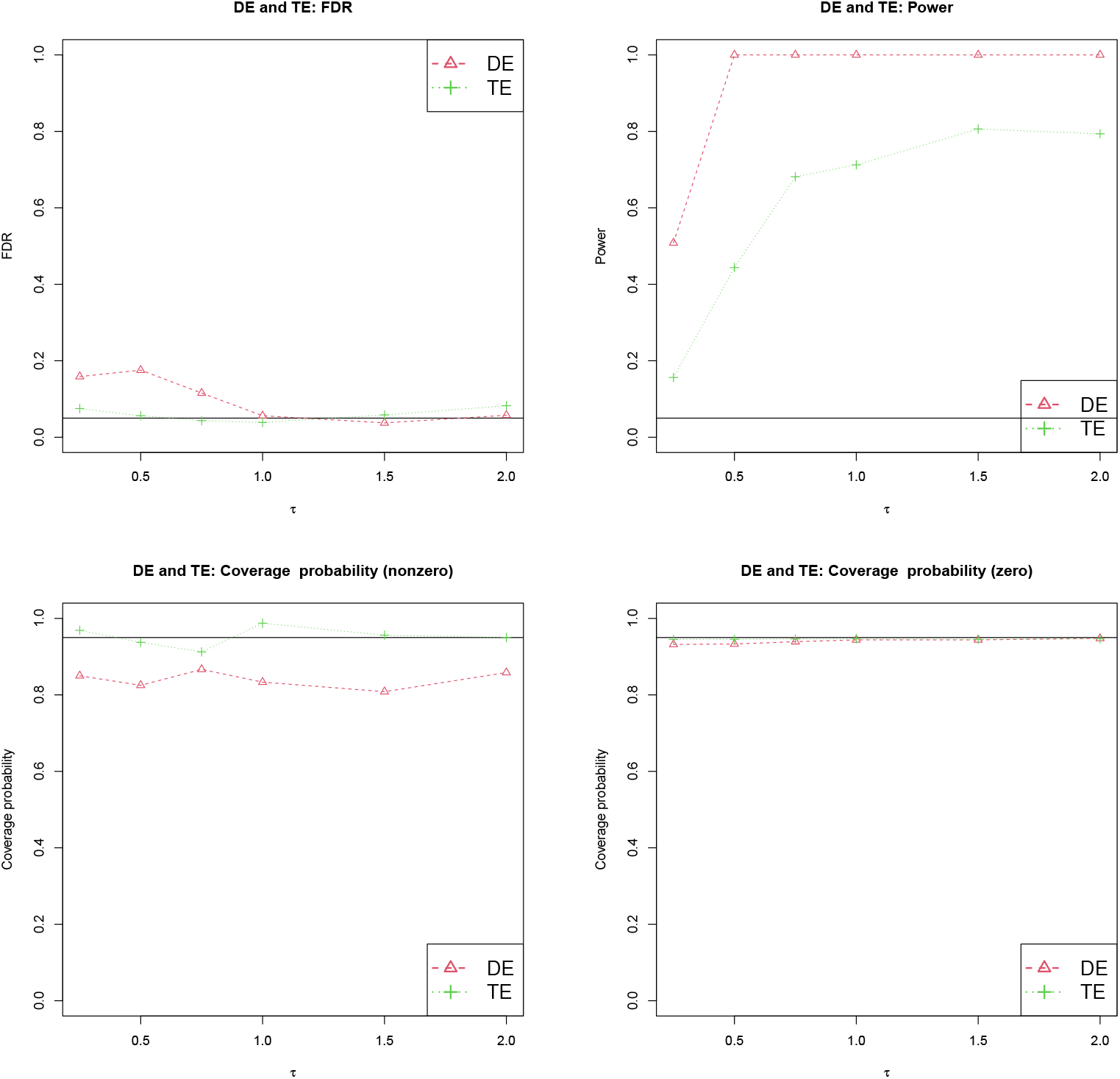
Complimentary inference of total effect (TE) and direct effect (DE) from MedDiC. Empirical FDR (upper left) and Power (upper right), and the marginal coverage probabilities for the true nonzero (lower left) and zero (lower right). The exposures are high dimensional(*q* = 400), and there are five true mediators among the *p* = 500 candidate mediators that are included in the model. The exposures are correlated (*r* = 0.4).

